# Structural variability and functional association in the epiphytic bacteria assemblies of freshwater macrophytes (*Myriophyllum spicatum*)

**DOI:** 10.1101/316893

**Authors:** Ling Xian, Tao Wan, Yu Cao, Junyao Sun, Ting Wu, Suting Zhao, Andrew Apudo Apudo, Wei Li, Fan Liu

## Abstract

The underlying principles influencing bacteria community assembly have long been of interest in the field of microbial ecology. Environmental heterogeneity is believed to be important in controlling the uniqueness and variability of communities. However, little is known about the influence of the host macrophytes on epiphytic bacteria assembly processe. Here, we produced two contrasting artificial water environments (eutrophic and oligotrophic) for reciprocal transplant experiment of *Myriophyllum spicatum*, to recover the colonization of epiphytic bacteria accompanied with plants growth. Comparative analysis addressed a higher species diversity in epiphytic bacteria than in bacterioplankton, and the highest microbiome richness in sediment. Our data revealed that the organization of epiphytic bacterial community was interfered by both plant status (i.e. branch number, net photosynthesis rate etc.) and water bodies (i.e. total phosphate, total nitrogen, pH etc.) while plant status effected the assembly in priority to water. 16S rRNA sequencing further indicated that the epiphytic assemblies were motivated by functionalization and interplay with hosts as a whole. The results complemented new evidences for the ‘lottery process’ in the epiphytic bacteria assembly traits and shed insights into the assembly patterns referring to functional adaptation across epiphytic bacteria and macrophytes.

**Importance:** A robust understanding of inter-adaptation between microbiome and the host plants have been established basing on vast majority of researches. However, great efforts were made mostly on rhizosphere microbiome. By contrast, referring to another representative group, macrophytes who composed of the freshwater ecosystem were relatively less investigated on such issue. Our study pioneered the experimental operation to interrogate the triadic relationship among macrophytes, epiphytic bacteria and water body. The research present here showed significant exemplar on discussion of plant associate bacteria adaptation taking account of host colonization as well as the epiphytes. The results expand the hypotheses of bacteria assembly principle and provides potential leads on understanding of plant - microbe interactions.

## Introduction

The relationship between bacterial composition and environment and the underlying principles of bacterial community assembly have triggered great debates in field of microbial ecology for decades (1–3). Two main principles have been control bacterial community structures: one is the ‘niche process’ that emphasizes the importance of biotic/abiotic factors, such as habitat environment in influencing the community structure of a species (4–6), the other one is a ‘neutral process’ that highlights the role of stochastic processes, to explain differences observed among communities in dispersal and specification (7–9). Recent investigations have suggested that these two mechanisms are interactive, rather than mutually exclusive, in their control of the diversity and composition of microbial communities in various ecosystems (3, 10). Among freshwater ecosystems, many studies have revealed the combined effects of niche and neutral processes on bacterial communities at both spatial and temporal scales (11–15). For instance, lacustrine bacterioplankton assembly could be driven by pH-related niche and neutral processes (14) and could also shape a unique biogeographical pattern under the two models (12).

In addition to bacterioplankton, there is another large community of bacterial in the form of epiphytes, that are believed to be important in modifying macrophyte growth and development. They not only could enhance nutrient cycling through nitrogen fixation (16), but also act as a source of carbon dioxide and organic compounds for the host plants (17). Beyond that, epiphytic bacteria can affect the photosynthetic activity of macrophytes through shading by forming thick biofilms on the plant surfaces (18). Despite this, our knowledge of the effects of epiphytic bacteria eon community ecology and dynamics (19), on is poorly known, especially the relations among bacteria, plants host and environment. This deficiency further impedes us to survey the underlying principles of their assembly formation on macrophytes. Unlike the other bacterial habitats within freshwater environments, macrophytes create a very different environment for epiphytes due to the high dispersal and propagation rates of plants through vegetative strategy (e.g. shoot dispersal) (20). This would continuously provide new opportunities for bacteria colonization which is particularly vigorous at the initial stages or when macrophytes colonize a new environment. On the other hand, since epiphytic bacteria colonization primarily originates in the aquatic environment, different environmental factors, like spatial and temporal ones, would probably influence their community structure and assembly, even with comparable niches in similar environments (9). Moreover, the heterogeneity of water will also introduce niche effects to macrophytes. For example, the eutrophic lakes have lower transparency and a higher potential for algal blooms compared to the oligotrophic lakes. Thus, macrophytes survival in eutrophic environments is generally impaired, while bacteria are likely to exist and compete for the carbon resources. Eutrophic aquatic ecosystems could also be toxic to macrophytes as a whole. All these make it extremely difficult to examine bacterial community structure, especially considering the characteristics of its initial assembly stages at natural environment (21). *Myriophyllum spicatum*, a submerged plant species from Haloragidaceae family which was widely distributed in freshwater habitats. It is noticeable that *M. spicatum* is reportedly highly tolerant to ammonium-nitrogen toxicity (23) which provide optimal opportunity to investigate the complex associations among epiphytic bacteria, host plants and different water bodies. Herewe selected two populations of *Myriophyllum spicatum*, as hosts from lakes with differentt trophic state. The sampling shoots were synchronized and reciprocally transplanted in the artificial freshwater ecosystems. With operative and successional observation, we characterized the composition of microbiome in different ecological niches including host and ambient compartments, to unravel the probable mechanisms on the assembly. Our work would provide more valuable perspectives on bacteria and macrophytes, taking the interaction with water condition into account. The efforts could also facilitate the integrative framework for ecological theory.

## Results

### Taxon and sequences information

A total of 2,591,428 effective tags were obtained from all 47 samples. The taxon tags and OTUs (97% identity) were both generated (Table S1). The number of OTUs per sample ranged from 812 to 5,717 with an average value of 2,799. Taxon information and the sequence numbers were generated according to the generated OTUs. There were 48,893 sequences (94.4%) at the class level and 185 classes were categorized. OTUs mainly comprised five classes including *Alphaproteobacteria* (30.6% ± 15%, based on OTU relative abundance), *Betaproteobacteria* (15.9% ± 10.7%), *Gammaproteobacteria* (9.3% ± 8.8%), *Planctomycetia* (5.7% ± 11.7%) and others (5.6% ± 5.4%)

### Bacterial diversity

Alpha diversity, the microbial diversity within each sample, was analyzed based on the OTU relative abundance and the Shannon index (Fig. 1). The results indicated no differences in diversity among the 4 treatments within each bacteria group. But for each treatment, the OTU richness was highly dependent on location interval with richness highest in the sediment intermediate in leaf and stem samples, and lowest in water samples. PCA analyses revealed strong clustering of bacterial communities according to the different space interval (stem, leaf, sediment, water) (Fig. S1). At the OTU level, PC1 explained 42.5% and PC2 13.9% of the total variation. The leaf and stem samples were clearly distinguished from sediment and water samples but did not cluster completely according to their respective plant compartment.

**FIG 1.**
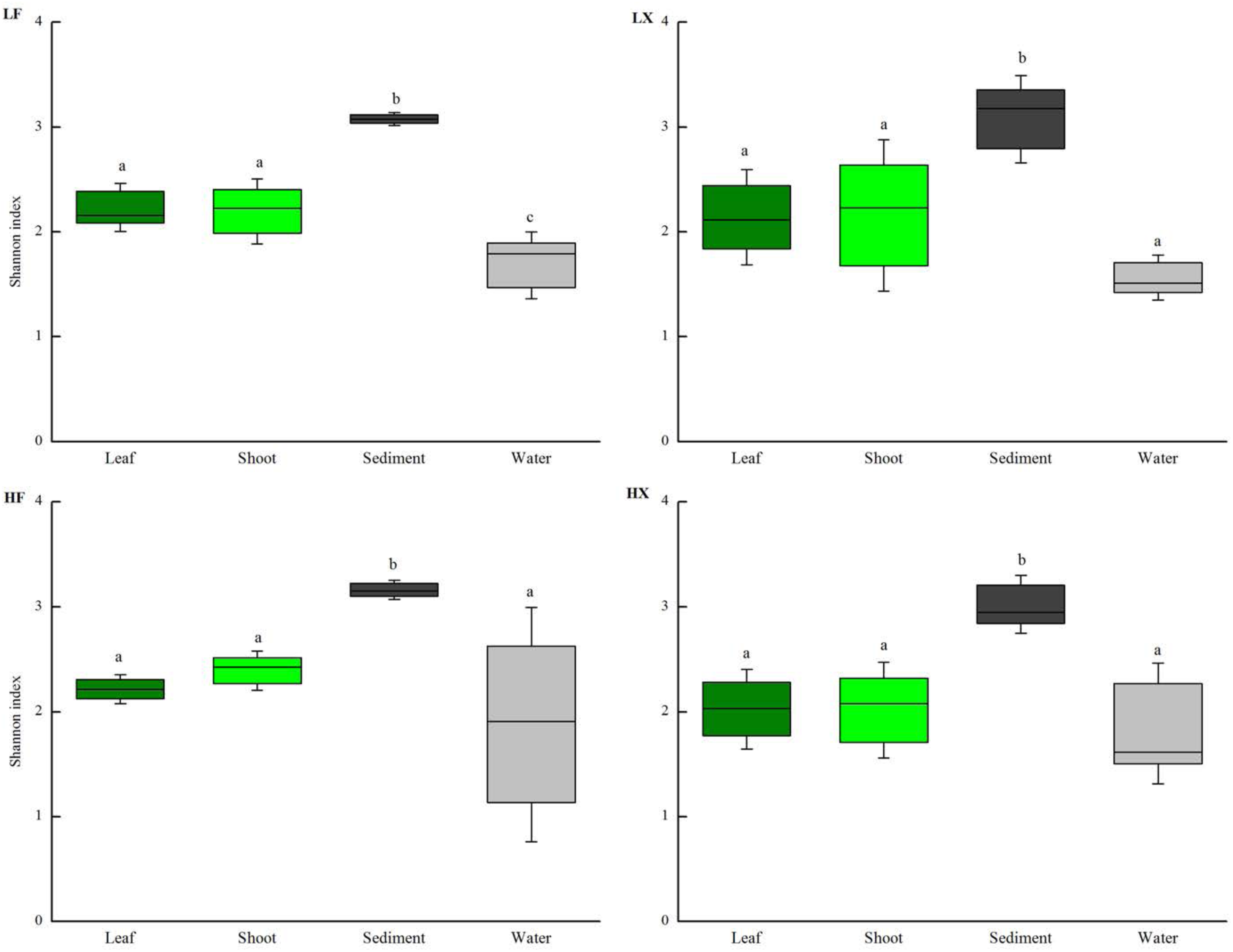
The Shannon diversity estimates of bacteria communities among leaf epiphytic bacteria, shoot epiphytic bacteria, sediment bacteria and water bacteria under four treatments (LF, LX, HF, HX). a, b, c shows the differences among different bacteria groups based on one way ANOVA.

### Members of the bacterial microbiome at different locations

Phylum distribution of the OTUs and relative sequence abundance of bacterial phyla associated with different ecological niches were shown in Fig. 2A. Phylogenetic composition of the community microbiota at the class and genus levels were examined, which differentiate the bacterial communities in the different space interval. According to the top 10 species comparison, *Alphaproteobacteria* is mostly abundant in epiphytic bacteria while *Betaproteobacteria* and *Gammaproteobacteria* were dominant in environments especially in the water and the *Deltaproteobacteria* waere rich in sediment. Besides the class species differentiation, analysis of the top 10 genera showed that *Erythromicrobium* and *Hyphomicrobium* belonging to *Alphaproteobacteria* were particularly dominant in epiphytic bacteria. *Hydrogenophaga*, *Massilia* and *Rheinheimera* belonging to *Betaproteobacteria* and *Gammaproteobacteria* were especially dominant in the water (Fig. 2B). Virtually all identified bacterial phyla displayed a significant space interval effect, regardless of initial treatments of plant hosts. Besides this, these are no composition differences in epiphytic bacteria related to nutrient states but the genus *Hydrogenophaga* was dominant in the high nutrient environment (Fig. 2B).

**FIG 2.**
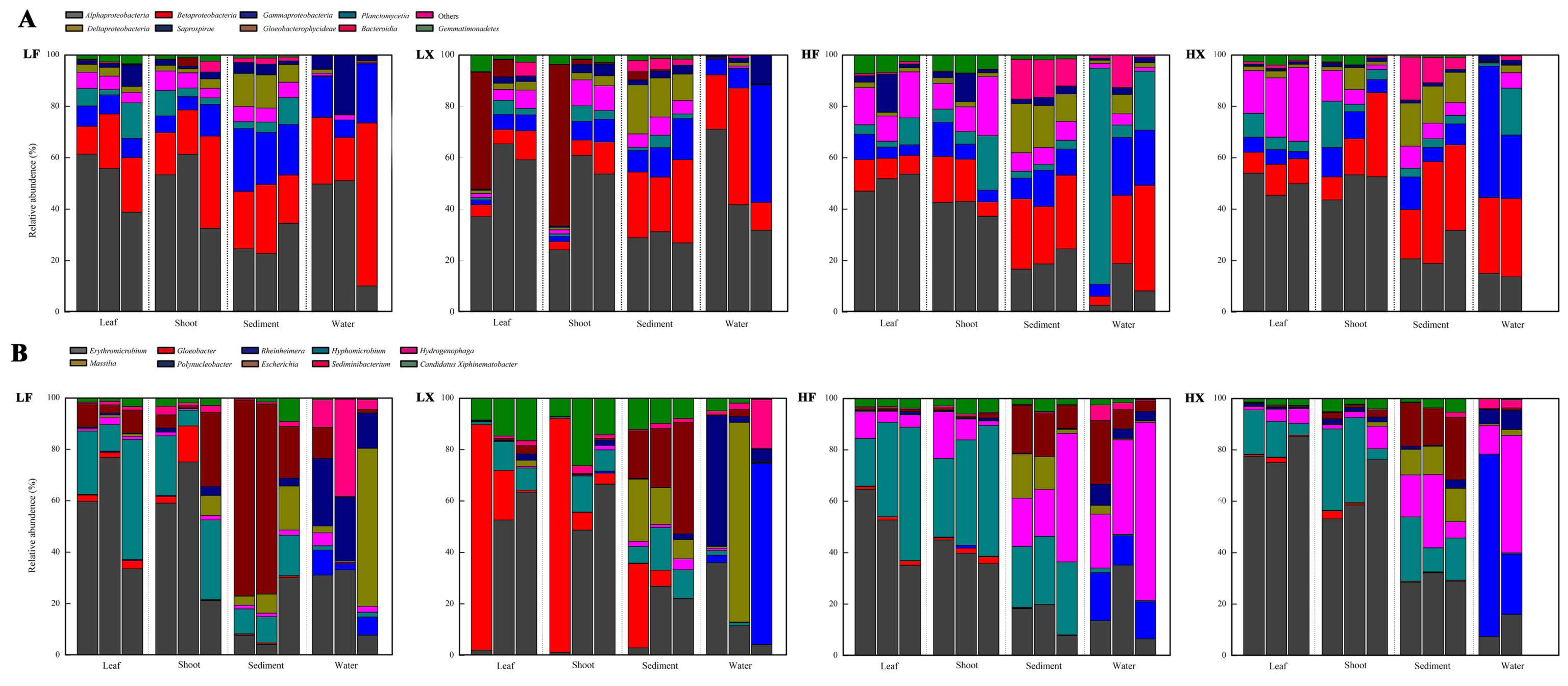
Top 10 distribution of the OTUs at class level (A) and top 10 identified distribution of the OTUs at genus level (B). The relative abundance associated with the leaf epiphytic bacteria, shoot epiphytic bacteria, sediment bacteria and water bacteria under four transplant treatments (LF, LX, HF, HX). The biological replicates (3 replicates for each except the water bacteria in HX treatment) are displayed in separate stacked bars.

### Relations among epiphytic bacteria, plant hosts and environments

The plants status and the physico-chemical data of the culture water system are presented in Table S2. For the plant, nearly all the morphological and physiological data were higher in the high nutrient site (HF and HX) except the Chla:b ratio and HCO_3_^-^ use. The Chla/b was higher in the low nutrient sites (LF and LX) while HCO3-remained consistent between the two nutrient states. For physico-chemical data, the temperature and Oxidation Reduction Potential (ORP) of the plant treatments were higher in Low nutrient site (LF and LX), while the parameters Conductivity (C), Salinity (SAL), Total Dissolved Solids (TDS), Total Phosphorus (TP), Total Nitrogen (TN) and Chlorophyll a (Chla) were higher in high nutrient site (HF and HX). The concentration of TP and TN in HF and HX were 40-and 15-times higher, respectively, than in LF and LX. However, the pH remained constant in all four treatments. Compared with plant status and environmental factors related to epiphytic bacteria, the plant status such as photosynthesis rate for leaf bacteria and the plant length for shoot bacteria were more related to the differences of BCC than environmental factors (Fig. 3).

**FIG 3.**
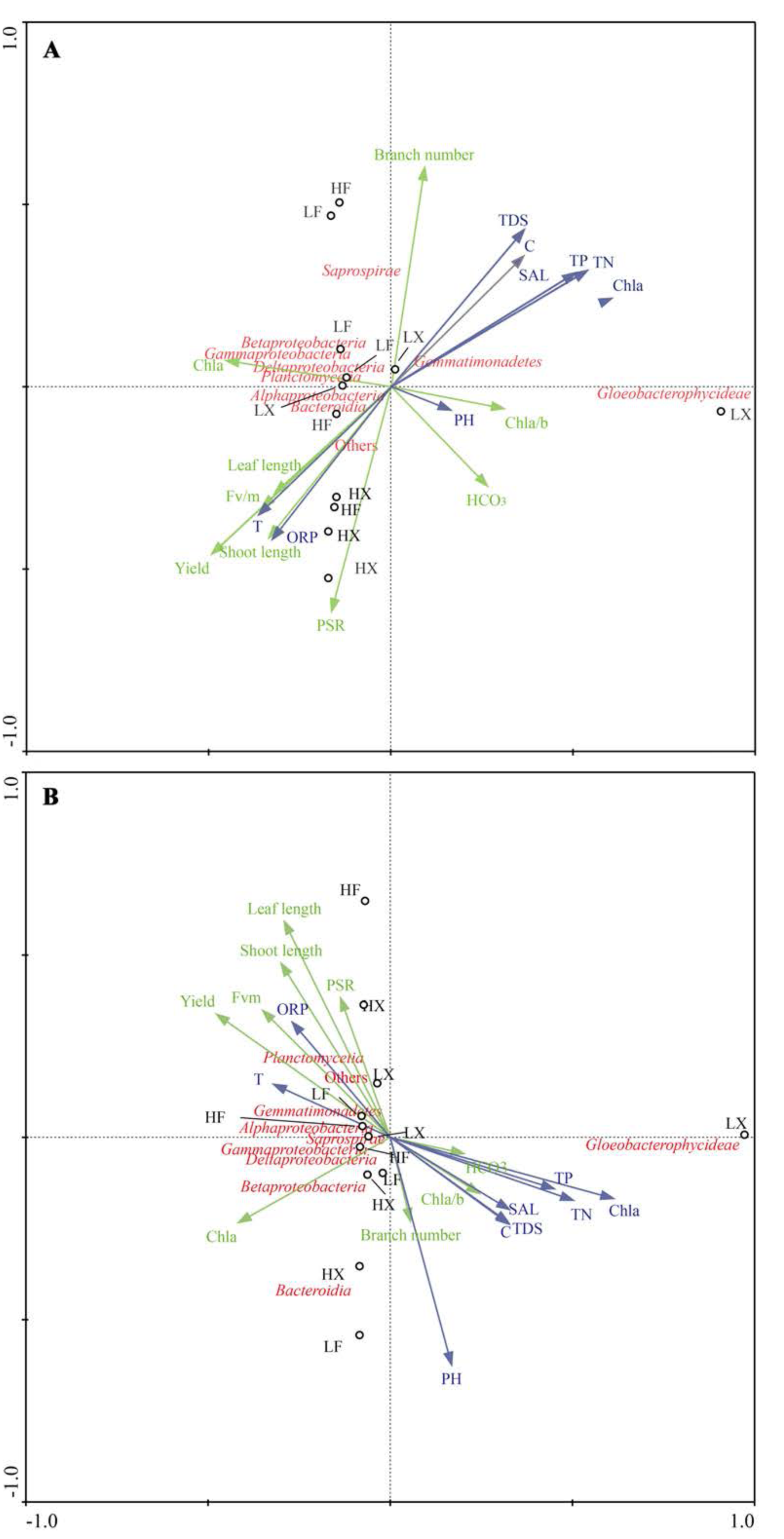
Canonical correspondence analysis (CCA) of the possible effect of macrophytes (green arrows) and environmental conditions (blue arrows) on distribution of the OTUs at class level (red color) associated with leaf epiphytic bacteria (A) and shoot epiphytic bacteria (B). LF, LX, HF, HX represent the four transplant treatments and three replicates for each treatment (black color). The macrophytes and environmental conditions are shown on Table S2. Macrophytes: PSR: Net photosynethsis rate, HCO_3_^-^: the ability to use HCO_3_^-^, Chla: Chlorophyll-a, Chla/b: Chlorophyll-a/ Chlorophyll-b, Fv/m: the quantum efficiency of open photosystem II centres, Yield: the real PSII photochemistry in a steady state of electron transport. Environmental water: T: Temperature, TDS: Total Dissolved Solids, C: Conductivity, SAL: Salinity, ORP: Oxidation reduction Potential, TP: Total Phosphorus, TN Total Nitrogen, Chla: Chlorophyll-a.

### Variation in functional association

In our study, the treatments based on the two different nutrient states including different transparency and nutrient level might cause the different use in light and nutrient. In order to compare functional differences among different bacteria and different treatments, The KEGG Orthologues (KOs) were chosen related to the enzymes in Calvin cycle for carbon fixation (ko00710): RuBisCO (K01601, K01602, K1807, K1808); GAPDH (K00134, K00150); PGK (K00927); PRK (K00855); TRK (K00615) and Aldolase (K01623, K01624, K02446, K03841, K04041, K11532). Enzymes related to ammonia assimilation were also analysed: glutamate synthase (GOGAT) (K00265, K00266); Fd glutamate synthase (Fd-GOGAT) (K00284), glutamine synthetase (GS) (K01915) and glutamate dehydrogenase (GDH) (K00260, K00261, K00262).

And globally, remarkable differences were observed between epiphytic and environmental bacteria. Nutrient treatments suggested to be a considerable issue in the function variation patterns. In detail, among all OTUs related to the Calvin cycle in carbon fixation, the OTUs of GAPDH were significantly higher for epiphytic bacteria and sediment bacteria in high nutrient treatment than those in lower but few differences for water bacteria (Fig. S2). Furthermore, among all the OTUs related to ammonia assimilation, the OTUs encoding glutamate dehydrogenase (GDH) also higher but glutamine synthetase (GS) lower for epiphytic bacteria in high nutrient treatment but the environment bacteria changed little (Fig. S3).

### Discussion

The leaves of *Myriophyllum spicatum* are pinnate and the leaflets needle-like, offering a large surface area for colonization by epiphytic bacteria compared to other macrophytes. Through simulating the natural macrophyte growth, we tried to characterize the composition of the epiphytic bacteria community and to unravel the relationship among the epiphytic bacteria community, plant hosts and environments as well as the potential community assemblages of the epiphytic bacteria.

### Diversity and community composition of macrophyte epiphytic bacteria

Increasing evidence has indicated a close relationship between epiphytic microbial communities and living hosts in freshwater and marine environments (37–39). He et al. (19) reported that the diversity of epiphytic communities on *Potamogeton crispus* and was higher than in the bacterioplankton. Our results are consistent with this point but also provided an in-depth assessment of the diversity profile of the bacteria communities. The high epiphytic diversity might be because the interface between water and plant hosts endows epibiotic bacteria with competitive advantages to acquire nutrients from both media, especially for carbon sources through the plants in productivity and element cycling (19, 40). Besides this, the community composition of bacteria showed that *Alphaproteobacteria* were dominant in epiphytic bacteria while *Betaproteobacteria* and *gammaproteobacteria* were dominant in the water. Although *Alphaproteobacteria* had diverse functions and few commonalities, the dominant genus in the *Alphaproteobacteria* in our study might indicate the functional role they played. For instance, recent studies indicate that the *Erythromicrobium* is likely to produce bacteriochlorophyll a and carotenoid for photosynthesis (41) while *Hyphomicrobium* prefers using carbon by the way of carbon dioxide (42). Furthermore, the *Betaproteobacteria* and *gammaproteobacteria* detected in water, together with the other top genera closely related, they might not be functionalized as the same as the *Alphaproteobacteria* in epiphytic bacteria. For example, *Massilia* is a phosphate solubilizing bacteria (43) while the *Hydrogenophaga* has great deal of degrading ability (44). These varying results for epiphytic and environmental bacteria indicate that the epiphytic bacteria assembled might be conferred to plants host through some specific functions (i.e. carbon use), while the environmental bacteria did not.

### Relationship between epiphytic bacteria and nutrient states

Besides the plants host, the nutrient states also affected the epiphytic bacteria composition together with the plants. Although the diversity analysis above indicated differences between the epiphytic and environmental bacteria, this difference is profound in low nutrient level. The epiphytic bacteria had a higher diversity in low nutrient environment. This may due to the plants status in different nutrient states (Table S2). The leaf area is thought to be crucial for epiphytic bacteria diversity, the larger leaf area always causes a higher bacteria diversity. In our study, we used leaf length instead of leaf area considering the nature of leaf morphology of *M. spicatum*. Interestingly, the longer leaf length in higher nutrient level caused lower bacteria diversity. We think this may due to our sampling strategy. Actually after we transplant the individuals, we found that the leaf length always higher but the branch number was lower in higher nutrient levels (Table S2). Besides that, when we sample the epiphytic bacteria one year after the start of the experiment, we used the same weight of the leaf, which further weakened the effect of total leaf area on the epiphytic bacteria composition.

More than that, our previous observation on the plants growth had revealed that the host individuals were likely to increase their shoot length to the surface, especially in higher nutrient water, to reach a balance of photosynthesis related status and the bacteria diversity (45). Our CCA analysis further supported this as the plant net photosynthesis rate (PSR), the yield value and the branch number were more related to epiphytic bacteria community composition compared to the other factors. The net photosynthesis rate is of ability for plants to use the carbon from environments while the yield value to transfer an electron to active one. These two factors may cause the different compostion in epiphytic bacteria. attributing to the competition for carbon sources under different nutrient state. The higher photosynthesis rate enables plants to use more carbon under light limitation conditions. This may cause the carbon source limitation for epiphytic bacteria, leading the low diversity of them. Our function prediction comparison on carbon fixation also give this evidence for the high abundance preference through the GAPDH (others have no differences) under high nutrient treatments for epiphytic bacteria for competition (Fig. S2).

Besides the plant status effects, the environmental factors such as water conditions, including nutrients (46), pH (14), salinity (47), temperature (48), and productivity (49) always proved to be the driving forces on the bacteria communities. PH was thought to be the main factor that affect the bacteria composition (14). Somehow, it was not well-supported in our study since no obvious differences on pH values within the two nutrient treatments. But we noticed the nitrogen level (TN) might introduce the differences in epiphytic bacteria from low to high nutrient treatments. Alternatively, this may be better understood with respect to function related to nitrogen metabolism (Fig. S3). In our study, the OTUs on glutamate dehydrogenase (GDH) are dominant under higher nutrient conditions (H nutrient sites) while glutamine synthetase (GS) and formamidase are dominant in low nutrient conditions. GDH catalyzes the reductive amination of 2-ketoglutarate by ammonia to give glutamate in an NADPH-dependent reaction (50). It had been reported to be an alternative way for ammonia assimilation, especially when the ammonia becomes toxic at high concentration (51). Higher levels of eutrophication imply higher ammonia toxicity which increased GDH production. Under high nutrient levels and associated lower transparency, plants tend to grow rapidly to the water surface to obtain light (45). In many organisms, the GS/GOGAT pathway facilitates the assimilation of ammonia present in the medium at concentrations lower than 0.1 mM (52). Consequently, the GS was crucial for ammonia assimilation through GS/GOGAT pathway, which was involved in the common pathways under ordinary nitrogen levels (52, 53).

### Macrophyte epiphytic bacteria underlying assembly principles

Niche and neutral models are the two primary theories that have been defined for bacteria assembly strategies (14). However, the results from the present study could not be sufficiently explained through either the niche or neutral models. Although the differences in ecological niches influence BCC, there was still considerable divergence in the same nutrient water system (e.g. differences in BCC between the epiphytic bacteria and the environmental samples within similar nutrient tanks). According to neutral theory, the species will occupy the space through random immigration, birth or death in a case where the trophic state is equal for all the species (8, 9). However, we observed striking differences among the samples even in the similar nutrient conditions (Table 1). Therefore, we suggest the ‘lottery hypothesis’ might be more appropriate to explain the epiphytic bacteria assembly patterns observed (39, 54, 55). The lottery model facilitates assembly for specific functions when bacteria colonize macrophyte surfaces. Within freshwater, the macrophytes responded to the stress as morphological and physiological changes (Fig. 4). For example, the lower transparency introduced by eutrophic water lead to an increase of plant shoot length, branch number, chlorophyll content etc. The epiphytic bacteria on the surface of the macrophyte leaves, will benefit from this by having more abundant source of carbon as well as light than water (56). Thus, the *Alphaproteobacteria* and related genus detected here are probable examples as photosynthesis and carbon metabolism exploiter. More than that, the water also has direct influences on both epiphytic and environmental bacteria (Fig. 4). It is noticeable that the epiphytic bacteria preferred to assimilate ammonia at the higher nutrient states despite their composition varied little, which indicated additional help on host plant resistance to nutrient toxicity (Fig. 3 and S3). Instead, the composition of environmental bacteria showed no specific and functional preference when responding to different water niches. Accordingly, suggest that there may be an interaction between niche and lottery effects, which is crucial for community assembly in macrophyte epiphytic bacteria (54), and the differences in BCC should be examined from a perspective of functional assembly instead of only species diversity (55). Although part of our conclusion is based on the predicted functions rather than metagenomics, they still offer a new explanation to understand the BCC at the species level. It would be expected that higher accuracy rate was calculated when metagenomics invovled (34) and the direct impact of epiphytic bacteria on macrophytes hosts will step forward in the near future.

**FIG 4.**
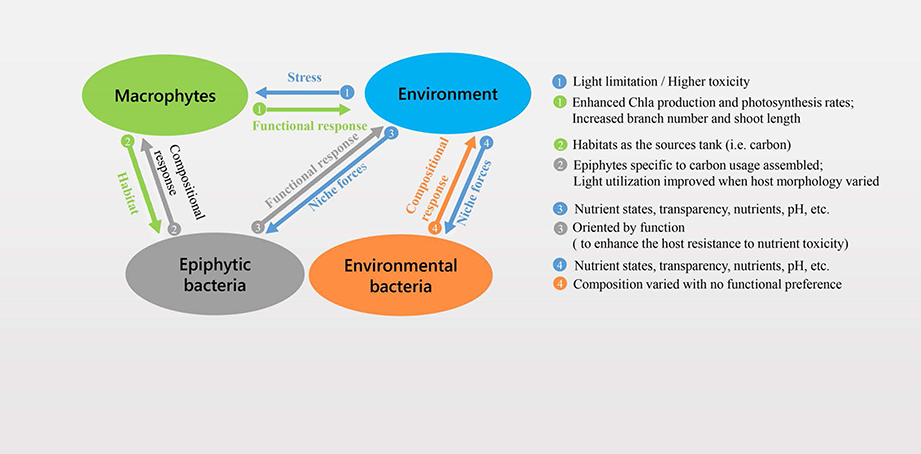
Summary of the triadic relationships among macrophyte hosts, bacteria (epiphytic bacteria and environment bacteria) and the environments.

## Material and methods

### Information for host plant and study sites

*Myriophyllum spicatum*, a submerged angiosperm belonging to the Haloragidaceae and is widely distributed in freshwater habitats, including eutrophic lakes (22). It is also reported for higher adaptive ability for nutrient toxicity (23). The experimental sites were on the shores of Fuxian and Xingyun lakes that are located in the center of the Yunnan plateau, South West China (Fig. 5A). The two lakes are connected by Gehe River (24, 25), Fuxian Lake is an oligotrophic lake (labelled as L, low nutrient condition), while Xingyun Lake is a eutrophic throughout the year (labelled as H, high nutrient condition). *M. spicatum* individuals distributing in these two lakes have been well-studied in Han et al., 2014, Apudo et al. 2016 (23, 24).

**FIG 5.**
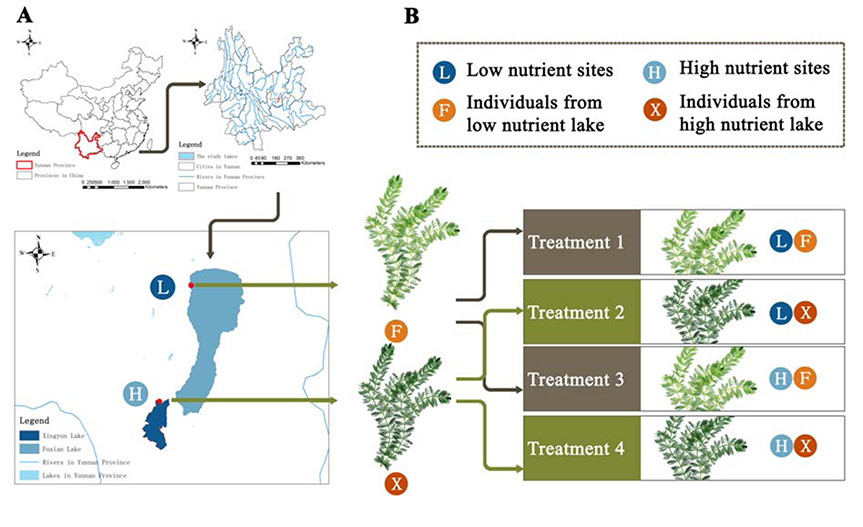
The study sites (A) and the reciprocal transplant treatments (B).

### Reciprocal transplant

Transplant experiments were conducted at two study sites adjacent to the lakes in order to investigate the combined effect of nutrition/trophic status and plant origin (Fig. 5B). In May 2014, 10 cm apices of *M*. *spicatum* were collected from Fuxian Lake (labelled F, for individuals from Fuxian Lake), and Xingyun Lake (labelled X, for individuals from Xingyun Lake). The apices were washed with sterile water and put into plastic pots (0.15 m deep and 0.26 m diameter) with sediment and water from L Lake. We repeated the same sampling process in H Lake site. Thus, four treatments (LF, LX, HF, HX) were generated. Each pot was placed into each cultivation tank (0.9 m deep by 0.45 m diameter) filled with water from L and H lakes Thus, each tank has one plant individual.

The water in all treatments was renewed every three days to maintain the homogeneity to the natural habitats. Water quality indices, including Temperature (T), pH, Total Dissolved Solids (TDS), Conductivity (C), Salinity, (SAL), Oxidation reduction Potential, Total Phosphorus (TP), Total Nitrogen (TN), and Chlorophyll-a (Chl-a) were measured in all four treatments. The plant morphological and physiological data including Branch number, Shoot length, Leaf length, Chlorophyll fluorescence (Fv/m and yield), Chlorophyll of leaf (Chla, Chla/b) and Photosynthesis related data (PSR: Photosynthesis rate, HCO_3_^-^: The use of HCO_3_^-^ of the leaves) were also recorded.

### Bacterial samples collection

After one-year culture, epiphytic bacteria were harvested from the host plant leaves (LFL, LXL, HFL, HXL) and shoots (LFSt, LXSt, HFSt, HXSt) in the four treatments. Environmental bacteria samples were also collected from the water column (LFW, LXW, HFW, HXW) and the sediment surface (LFS, LXS, HFS, HXS). There were total four plants’ treatments, two epiphytic bacteria sources, and two classes of environmental bacteria. Three replicates were sampled from each treatment (48 samples) for further DNA extraction. Fresh leaves and shoots (0.5 g FW) from the same plant parts (15 to 5 cm from the top) were put in 50 ml sterile glass bottles with cleaning buffer (2 mM phosphate buffer solution, 0.01%v/v Tween 80) for epiphytic bacteria sampling. The samples were ultra-sonicated for 5 minutes (19). 400 ml water samples were collected from each tank at a depth of 0.5 m using sterile polyethylene bottles. Afterwards, the 50 ml bacterial suspensions of epiphytic bacteria and the 400 ml water samples were filtered using DURAPORE ^®^ Membrane filters (0.22 μm, Millipore, Ireland Rev). The filters were stored at -20 °C for later extraction of DNA. 0.25 g of the surface sediment was collected and stored at -20 °C.

### DNA extraction, amplification and sequencing

Epiphytic bacteria and water sample DNA were extracted by PowerWater^®^ DNA Isolation Kit (MOBIO Laboratories, Inc., Carlsbad, CA). Sediment sample DNA was extracted by PowerSoil^®^ DNA Isolation Kit (MOBIO Laboratories, Inc., Carlsbad, CA). DNA was diluted to 1 ng μL^-1^ and 16S rRNA gene of distinct region V4 was amplified using a specific primer 515F (GTGCCAGCMGCCGCGGTAA) - 806R (GGACTACHVGGGTWTCTAAT). All PCR reactions were carried OTU with Phusion^®^ High-Fidelity PCR Master Mix (New England Biolabs) under the following conditions: 25 cycles of denaturation at 94°C for 30 s, annealing at 55°C for 30 s, and extension at 72°C for 30 s, with a final extension at 72°C for 5 min. PCR products were run through electrophoresis on 2% agarose gel for detection. Samples with bright main strip between 400-450bp were chosen and purified with Qiagen Gel Extraction Kit (Qiagen, Germany). Afterwards, 47 qualified samples (one water sample from HX was lost) were sent to Novogene Bioinformatics Institution (Beijing, China) to be sequenced on an IlluminaHiSeq 2500 platform. The sequence data from the bacteria are submitted and available in Genebank (No. SRP107261).

### Sequence data analysis

Paired-end reads without barcodes and primer sequences were merged using FLASH (26) to generate the raw tags. The raw tags were filtered to obtain the high-quality clean tags using QIIME to control quality process (27, 28). The tags were compared with the reference database using UCHIME algorithm (29) and the effective tags were finally obtained after comparing reference database under UCHIME algorithm (29). Operational taxonomic units (OTUs) (97% identity) were retrieved using effective tags by Uparse software (30). Representative OTU sequences were aligned with the rDNA database (Greengenes) for OTUs taxonomic assignment using the MUSCLE software (31). The basic information about OTUs from all samples were calculated at the class level.

### Diversity analysis

OTU abundance information was normalized using a standard of sequence number corresponding to the sample with the least sequences. The relative abundance in each group was calculated according to the OTU abundance information. The alpha diversity of each group including Observed species and Shannon Index were calculated using QIIME software (28). The top 10 species at class and at the genus level were chosen to assess diversity among the four treatments and different bacteria. In addition, Principal Component Analysis (PCA) was used to compare the differences in bacterial community composition (BCC) among the all treatments by PC-ORD 5.0 (32). Canonical Correspondence Analysis (CCA) was used to detect the relationship between BCC, the plants status and the environmental factors for bacteria to identify the crucial factors for BCC. CCA was completed by applying Canoco for windows software (version 4.5) (33).

### Functional content prediction

We used 16S rRNA to predict metagenomic functional content in PICRUSt software (34). PICRUSt uses an extended ancestral state reconstruction algorithm to predict which gene families are present, and then combines gene families to estimate the composite metagenome (34). The OTU table was normalized by the “normalize_by_copy_number. py” module in PICRUSt, and the assigned Greengenes Ids to the Kyoto Encyclopedia of Genes and Genomes (KEGG) Orthology (KO) (35, 36). Datasets were used to predict the metagenome function by the “predict_metagenomes.py” module in PICRUSt. The contributions of OTUs to particular functions calculated using “metagenome_contributions.py.” script in PICRUSt. Afterwards, the thousand predicted functions acquired were collapsed into higher categories for further analysis.

## Acknowledgements

We thank Total Genomics Solution (TGS) Institute (Shenzhen, China) for function prediction analysis, Dr Brigitte Gontero and Dr Stephen Maberly for their kind help in checking the language. This work is supported by the National Key R & D Program of China (2016YFA0601001) and the National Natural Science Foundation of China (grant Nos. 31370262, 31670368, 31670369 and 31500296).

The authors declare no competing financial interests in relation to the work.

---

**FIG S1** Principal component analysis (PCA) between epiphytic bacteria and environmental bacteria (sediment bacteria and water bacteria) based on the distribution of the OTUs at the class level of operational taxonomic units (OTUs). HFL, HXL, LFL, LXL associated with leaf epiphytic bacteria, the HFSt, HXSt, LFSt, LXSt associated with shoot epiphytic bacteria, the HFS, HXS, LFS, LXS associated with sediment bacteria and HFW, HXW, LFW, LXW associated with water bacteria.

**FIG S2** Comparisons of contributions of OTUs to carbon fixation (represented by Glyceraldehyde 3-phosphate dehydrogenase). The relative abundance associated with the leaf epiphytic bacteria, shoot epiphytic bacteria, sediment bacteria and water bacteria under four transplant treatments (LF, LX, HF, HX). a, b shows the differences of KO abudance among different bacteria groups based on one way ANOVA

**FIG S3** Comparisons of contributions of OTUs to ammonia assimilation (represented by glutamate synthase, Fd dependent glutamate synthase, glutamine synthetase and glutamate dehydrogenase). The relative abundance associated with the leaf epiphytic bacteria, shoot epiphytic bacteria, sediment bacteria and water bacteria under four transplant treatments (LF, LX, HF, HX). a, b, c showed the differences of KO abudance among different bacteria groups based on one way ANOVA

